# High-Quality Nuclei Isolation from Postmortem Human Heart Muscle Tissues for Single-Cell Studies

**DOI:** 10.1101/2023.02.05.526322

**Authors:** Sarah Araten, Ronald Mathieu, Anushka Jetly, Hoon Shin, Nazia Hilal, Bo Zhang, Katherine Morillo, Deepa Nandan, Indu Sivankutty, Ming Hui Chen, Sangita Choudhury

## Abstract

Single-cell approaches have become an increasingly popular way of understanding the genetic factors behind disease. Isolation of DNA and RNA from human tissues is necessary to analyze multi-omic data sets, providing information on the single-cell genome, transcriptome, and epigenome. Here, we isolated high-quality single-nuclei from postmortem human heart tissues for DNA and RNA analysis. Postmortem human tissues were obtained from 106 individuals, 33 with a history of myocardial disease, diabetes, or smoking, and 73 controls without heart disease. We demonstrated that the Qiagen EZ1 instrument and kit consistently isolated genomic DNA of high yield, which can be used for checking DNA quality before conducting single-cell experiments. Here, we provide a method for single-nuclei isolation from cardiac tissue, otherwise known as the SoNIC method, which allows for the isolation of single cardiomyocyte nuclei from postmortem tissue by nuclear ploidy status. We also provide a detailed quality control measure for single-nuclei whole genome amplification and a pre-amplification method for confirming genomic integrity.

## Introduction

The field of cardiovascular disease previously has been dominated by studies that focus on the inherited genetic origins of disease, which have been accomplished by analyzing DNA that is typically isolated from easily accessible whole blood and sequenced with the use of whole exome sequencing or targeted gene panels^1–4^. However, these methods mask mutations with low variant allele frequencies, and new high-throughput sequencing methods and single-cell muti-omics studies can uncover the impact of somatic mutations in cardiovascular disease^5^. These recent technological developments have led to increased capability, and the accessibility of information obtainable from a single biological sample has been expanded^6,7^. Omics technologies, such as genomics, epigenomics, and transcriptomics, are increasingly employed in parallel to construct a more comprehensive picture of development and disease^8,9^.

Accompanying the rise in multi-omic studies is an increased demand for efficient single-nuclei isolation for high-yield, high-quality biomolecules, with sample quality well established for downstream analysis leading to the detection of biologically meaningful signals. To perform these studies, the tissue collection methods need revision along with the methodology for assessing tissue quality and establishing quality thresholds^10^. Furthermore, fresh tissue is rarely obtainable for the non-diseased human heart due to its inaccessibility and unavailability for biopsy. Non-diseased tissues are mainly collected from postmortem autopsies. The postmortem tissue quality has been thought to be impacted by the postmortem interval field (PMI)^11^, defined as the time between donor death and tissue preservation. These postmortem tissues are crucial in the biomedical field for the implementation of essential laboratory findings on human heart diseases and aging^10^. This is particularly pertinent in the case of postmortem human tissues where sample availability is limited, and tissue heterogeneity may mask associations between diseased and non-diseased tissue. Our laboratory’s primary focus is on cardiac aging, and we found that the NIH NeuroBioBank had availability of non-diseased human heart tissues.

However, many of the tissues did not have an RNA integrity number (RIN) or DNA integrity number (DIN) that was specific to the heart or a documented postmortem interval (PMI). This led us to investigate the tissue quality collected from the biobank. In this study, we provide a detailed strategy to select superior-quality heart tissue and a method for simultaneous isolation of single-nuclei DNA and RNA from human postmortem tissue, with resultant nucleotides meeting the high-yield and high-quality requirements for whole genome amplification and omics analyses. Notably, we determined that our Single-Nuclei Isolation from Cardiac tissue (SoNIC) methodology was essential to extracting high-quality single-nuclei, DNA, and RNA from heart muscle tissue. We report comprehensive details of nucleic acid yield and quality and consider the utility of covariates. We have established tissue quality determination markers from the collected human heart tissue cohort, recorded their PMI values, and analyzed them for association of RNA and DNA quality. We demonstrate through single-nuclei whole genome sequencing (WGS) and RNA sequencing (RNA-seq) based gene expression profile analyses that our protocol for isolating DNA and RNA from single-nuclei from postmortem tissue is appropriate for WGS, RNA-seq, and multi-omic studies.

## Methods

### Human tissue collection and sample preparation

All human tissues have been obtained from the NIH NeuroBioBank at the University of Maryland. Samples were processed according to a standardized protocol under the supervision of the NIH NeuroBioBank ethics guidelines. This study was approved by the Boston Children’s Hospital Institutional Review Board (IRB, S07-02-0087). Tissue from 106 humans formed our cohort: 33 from individuals with a history of myocardial disease, diabetes, or smoking, and 73 controls without cardiovascular disease. Healthy cases had no previous history of cardiovascular disorder, and the cause of death was unrelated to any cardiovascular disease condition. Pathological assessment of healthy subjects reported no disease-related pathology beyond that expected in aged individuals, and clinical diagnosis of disease cases were confirmed through medical records. Diagnosis, age, sex, and postmortem interval (PMI) of the specimens utilized are detailed in Table 1.

**Table 1a.**
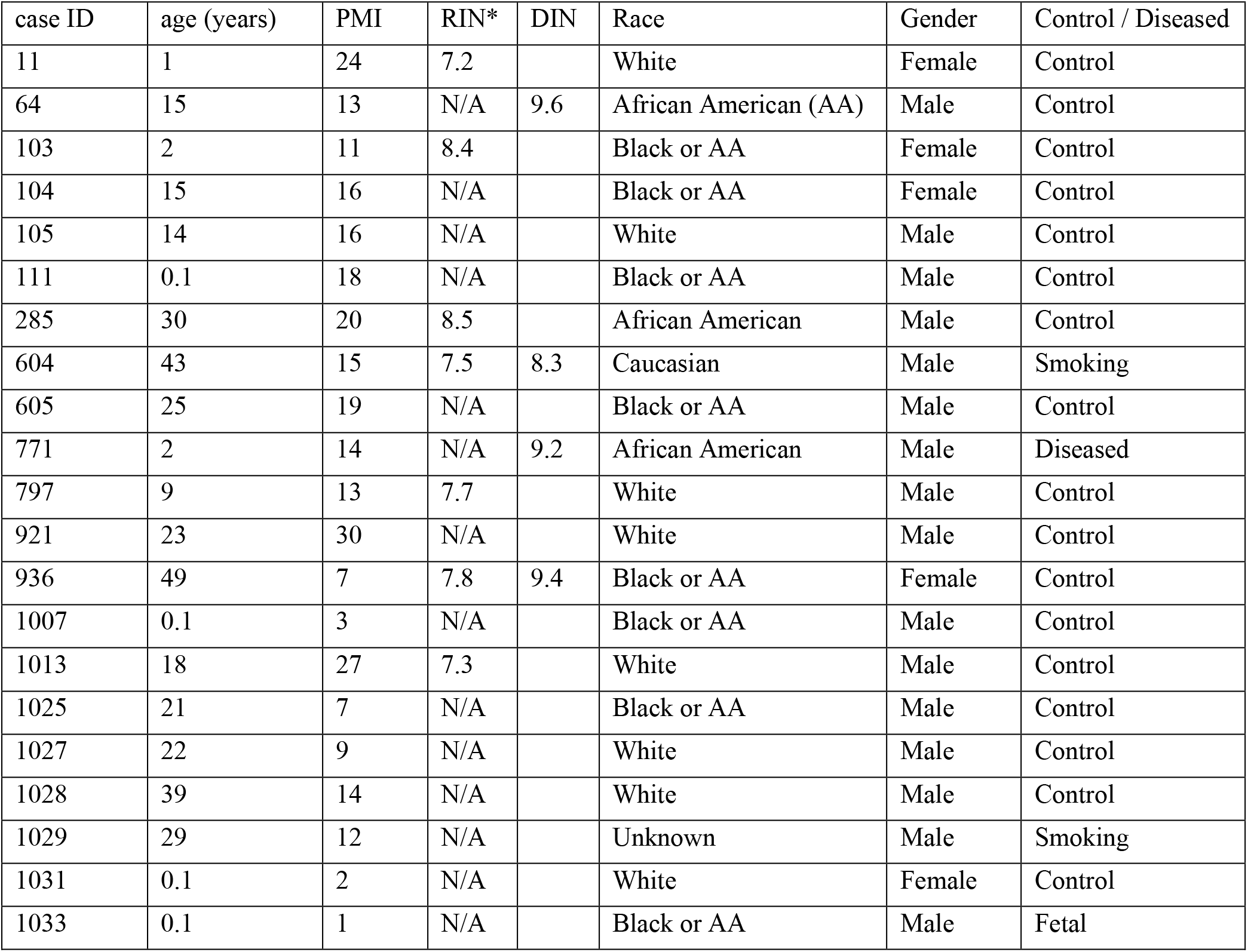
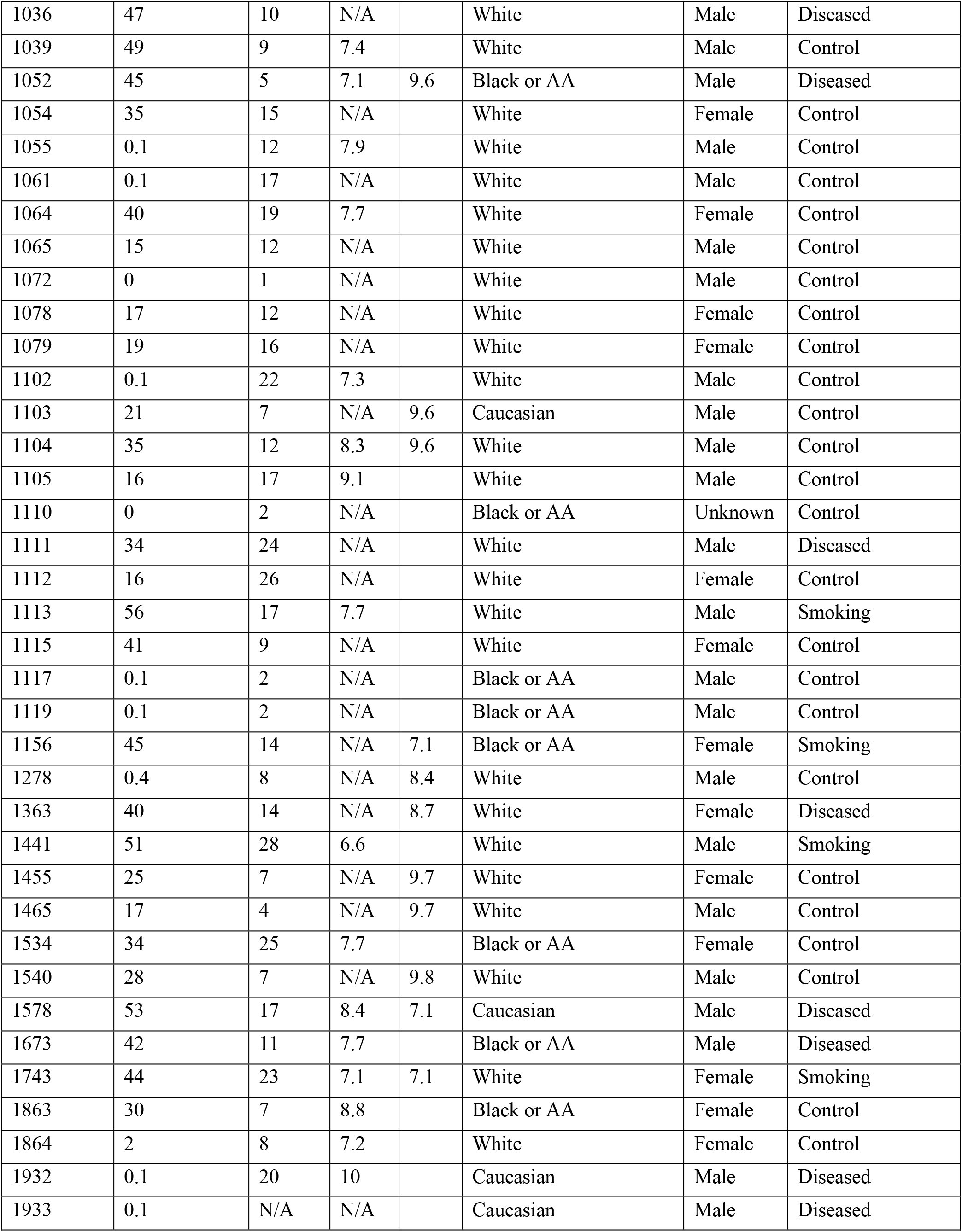
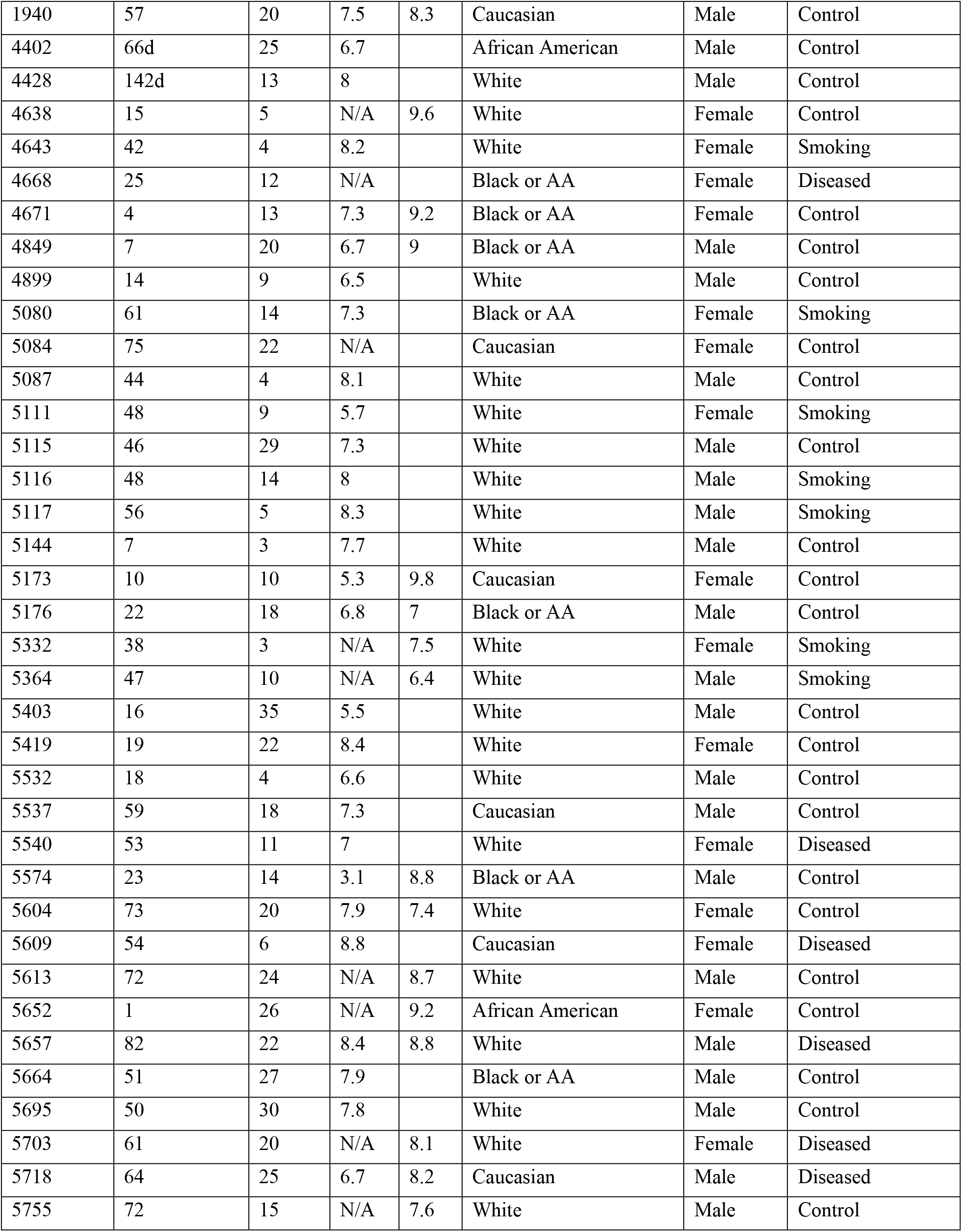
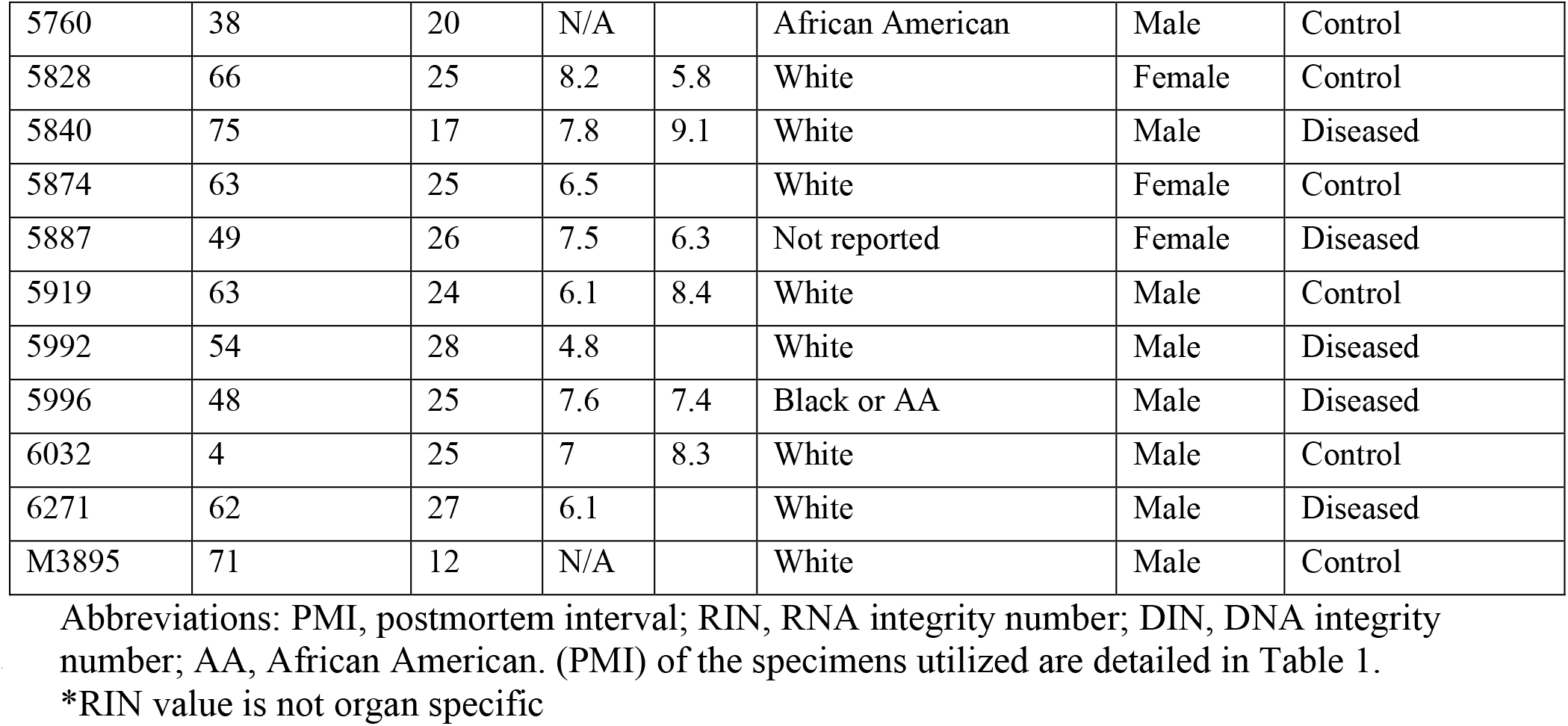
Characteristics of Heart Tissue Samples (n=106)

**Table 1b.** Characteristics of Tissue from Other Organs

| case ID | age (years) | PMI | RIN* | DIN | Organ |
| --- | --- | --- | --- | --- | --- |
| 604 | 43 | 15 | 7.5 | 8.3 | Heart |
| 604 | 43 | 15 | 7.5 | 8.5 | Liver |
| 604 | 43 | 15 | 7.5 | 9.0 | Kidney |
| 936 | 49 | 7 | 7.8 | 9.4 | Heart |
| 936 | 49 | 7 | 7.8 | 7.9 | Liver |
| 936 | 49 | 7 | 7.8 | 8.9 | Kidney |
| 1278 | 0.4 | 8 | N/A | 8.4 | Heart |
| 1278 | 0.4 | 8 | N/A | 5.9 | Liver |
| 5657 | 82 | 22 | 8.4 | 8.8 | Heart |
| 5657 | 82 | 22 | 8.4 | 6.3 | Liver |

\*RIN value is not organ specific

### DNA Extraction

Upon receipt of the tissue from the NeuroBioBank, DNA degradation was evaluated by isolating DNA from the tissue using the Qiagen EZ1 Advanced XL machine (Qiagen Catalog No. 9001875) and EZ1 DNA Tissue Kit (Qiagen Catalog No. 953034). Briefly, tissue was dissected with a scalpel on a cold block to produce approximately 34 mg of tissue. Then, 190μl of G2 buffer (Qiagen Catalog No. 1014636) and 10μl Proteinase K solution (Qiagen Material No. 101406) was added to the tissue. These samples were then vortexed for 15 seconds, briefly centrifuged for 15 seconds at maximum speed on the benchtop centrifuge and placed onto the thermo-mixer at 56°C until the tissue was dissolved (approximately 2 hours). Subsequently, all samples were pipette mixed, briefly centrifuged for 15 seconds at maximum speed on the benchtop centrifuge, and loaded onto the Qiagen EZ1 machine for extraction, with use of the EZ1 DNA Reagent Cartridge for tissue (from Qiagen Catalog No. 953034). The elution value was set as 200 μl for each of the extraction methods, and once extracted, all samples were stored at −20°C until quantification was carried out. Genomic DNA Screen Tape (Agilent Catalog No. 5067-5365) was used to determine DNA length. Tissues with fragmented DNA were not selected for further studies.

### Single-Nuclei Isolation from Cardiac tissue (SoNIC) methodology

To isolate single cardiomyocyte nuclei from frozen postmortem human tissue, we modified the protocol from Bergmann et al.^12,13^, as we started with a very small tissue sample. Briefly, 100 mg of tissue was dissected from the left ventricle of the heart (Figure 1a) with a scalpel on a cold block (Figure 1b) and resuspended <u>on ice</u> in 1 ml of lysis buffer containing 0.32 M Sucrose (VWR Catalog No. 97061-432), 10 mM Tris-HCl pH 8 (Invitrogen REF 15568-025), 5 mM Calcium Chloride (Sigma-Aldrich 21115-100ML), 5 mM Magnesium Acetate (Sigma-Aldrich 63052-100ML), 2 mM EDTA (Invitrogen REF AM9260G), 0.5 mM EGTA, and 1 mM DTT (Promega REF V3151). Resuspended tissue was transferred to a 5 ml Eppendorf tube (Fisher Catalog No. 14-282-301) and homogenized with a mechanical dissociator with use of reusable tips (Omni 30750H) at 24,000 rpm with 2 × 30 sec pulses per sample (Figure 1c).

**Figure 1.**
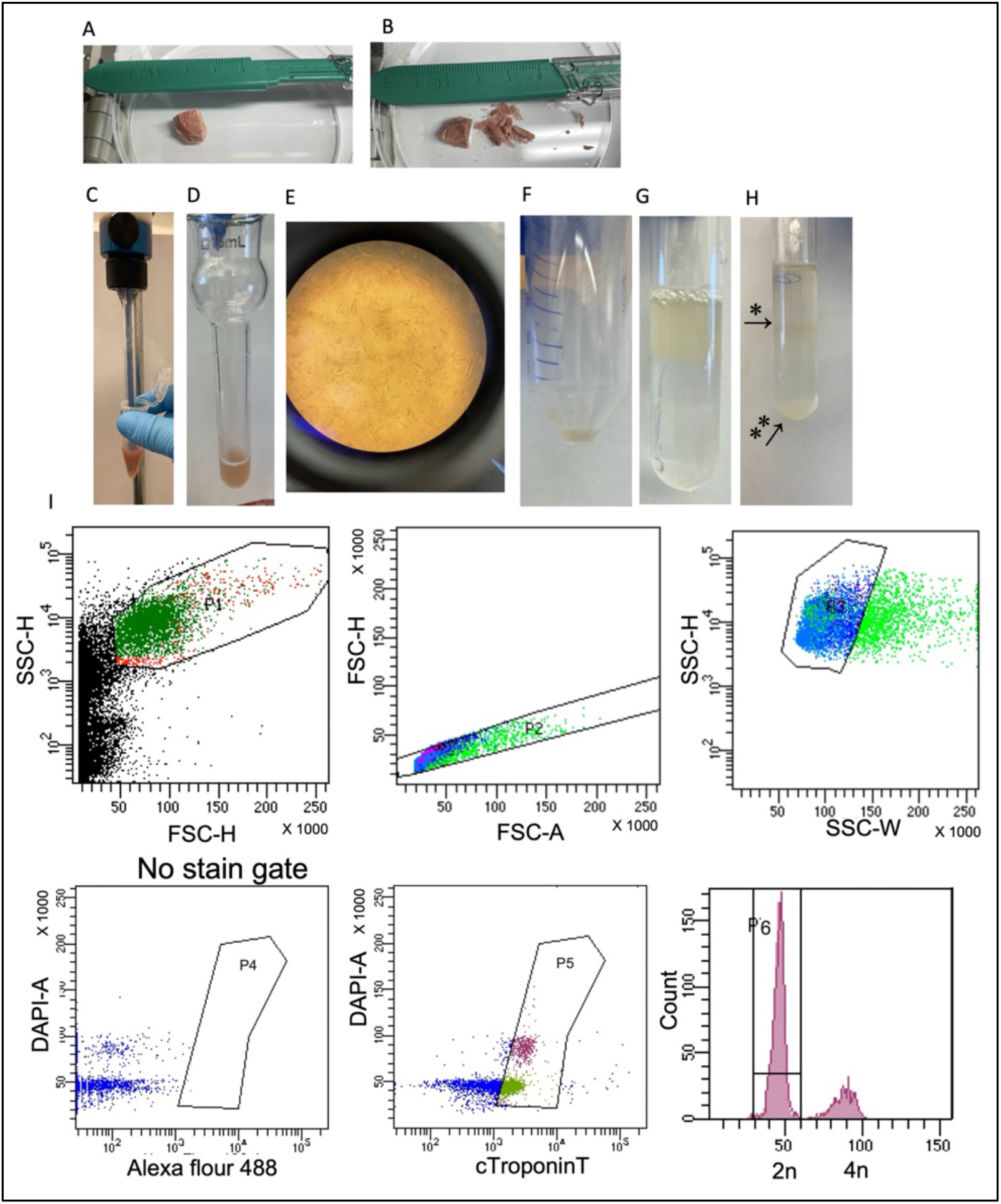
SoNIC method for the isolation of single-cell cardiomyocyte nuclei for use in downstream DNA and RNA analysis. (a) Tissue sample from the left ventricle of the heart (b) was dissected using a scalpel and transferred into lysis buffer for (c) mechanical dissociation and then transferred to (d) a douncer to break the cardiomyocytes (e) and free the nuclei. The sample was filtered and spun at 700xg for 10 minutes (f) and then the pellet was resuspended in lysis buffer and (g) laid on top of 5 mL of a sucrose buffer and then centrifuged at 13,000xg for 40 minutes (h) at which point a debris layer formed (*) and nuclei were pelleted on the bottom (**). The bottom pellet was resuspended in nuclei storage buffer and stained using a Cardiac Troponin T antibody and DAPI, after which the sample was filtered through the top of a falcon tube and sorted. (i) Cells were sorted based on Cardiac Troponin T (cTroponinT) status and/or DAPI intensity.

Tissue lysate was further diluted on ice with 2 ml of lysis buffer and dounced by hand with a 15 ml glass douncer and B pestle (Kontes Catalog No. 885300-0015) for 40-60 strokes (Figure 1d) to break the cardiomyocytes (Figure 1e) and free the nuclei. The crude lysate was consecutively filtered through 100 μm and 70 μm cell strainers (Corning Catalog No. 431752 and 431752). The filtered lysate was centrifuged at 700xg for 10 minutes at 4°C. In advance of the next step, Ultra-Clear Ultracentrifuge Tubes (Beckman Coulter Catalog No. 344061) were pre-coated with 10 mL of 1% BSA (Sigma Catalog No. A7409-50ML) in PBS (Life Technologies Catalog No. 10010072), inverted once, and stored for 40 minutes on ice, after which the solution was removed. The tubes were filled with 5 ml of a sucrose buffer containing 2.1 M Sucrose, 10 mM Tris-HCl pH 8, 5 mM Magnesium Acetate, and 1 mM DTT and the tubes were saved for use in the next step. After centrifugation, the supernatant was removed. The pellet (Figure 1f) was resuspended in 1 ml lysis buffer, added on top of the sucrose buffer (Figure 1g), and centrifuged at 13,000xg for 40 minutes at 4°C. After centrifugation (Figure 1h), the supernatant was removed and the nuclei were resuspended in Nuclei Storage Buffer containing 10 mM Tris-HCl pH 8, 70 mM KCl, 10 mM Magnesium Chloride (Sigma-Aldrich M1028-100ML), and 1.5 mM spermine (Sigma-Aldrich S3256-5G). Isolated nuclei were first stained for 40 minutes at 4°C with Cardiac Troponin T (Novus Biological NB120-10214AF488) at a 1:100 dilution in 0.01% BSA, and then stained with DAPI (Thermo Scientific REF 62248) at a 1:100 dilution for 5 minutes. Nuclei were strained through the 35 μm filter on the Falcon round bottom 5 ml polystyrene test tubes (Corning Catalog No. 08-771-23) and sorted with use of the fluorescence activated cell sorter FACSAriaIII (20 psi, 100-μm nozzle, Becton Dickenson Biosciences). The identity of heart muscle nuclei was determined by single-cell RNAseq and the quality of single-nuclei DNA was measured by multiplex polymeric chain reaction (PCR) and median absolute pairwise difference (MAPD) score after whole genome amplification.

### Evaluation of heart cell markers from single-nuclei RNAseq

The quality of heart tissue was also measured by single-nuclei RNAseq. Single heart nuclei were isolated using the SoNIC method as described. Each sample was transferred into a 5 ml polystyrene flow cytometry tube and sorted with the use of the 100-μM nozzle on the FACSAria III Cell Sorter (BD Biosciences Inc., Franklin Lakes, NJ, USA). The sorting strategy included doublet discrimination and selection of intact nuclei by sub-gating on DAPI-positive nuclei (Figure 1i). Doublet exclusion was performed by plotting the area for forward scatter (FSC) and side scatter (SSC) against the height (H) or width (W) or area (A); H versus W or A allows the separation of doublets from single-nuclei that are tetraploid and therefore contain more, 4n, amounts of DNA. Tetraploid nuclei containing 4n amounts of DNA have double the A and H values, whereas W is roughly the same as diploid cells containing 2n amounts of DNA. DAPI-positive nuclei were chosen based on ploidy (2n or 4n) and were sorted directly into a PCR tube containing a master mix as suggested by the 10X user guide. 3’ single-nuclei libraries were generated using the 10X Genomics Chromium Controller and following the manufacturer’s protocol for 3’ V3.1 chemistry with NextGEM Chip G reagents (10X Genomics Inc., Pleasanton, CA, USA). Final library quality was assessed with use of the Tapestation 4200 (Agilent Inc., Santa Clara, CA USA).

### Cell-type identification from 10X RNAseq data

After sequencing, the resulting sample FASTQ files from all samples were processed using CellRanger (v2.1.1) and Seurat package (v3.1.5) pipeline. Using the “mkfastq” and “count” commands we generate raw gene-barcode matrices and align them to GRCh38 Ensembl (v1.2.0). Combining multiple strategies, we compile a list of genes that are expressed in each type of cell in a specific way. By categorizing each nucleus as either coming from the type of target cell or not, we first estimated an AUC at the nuclei level and then predicted this class using the normalized expression of each gene. Using the edgeR function^14,15^ (filterByExpr (group=cell type), we eliminated genes whose counts were too low for testing. To determine marker genes, we selected protein-coding genes that were expressed in at least 25% of nuclei from the target cell type, with AUC for the target cell type greater than 0.60, a log-fold change, and an FDR adjusted P-value < 0.01 ^16^. The cell-type labels for each cluster were assigned based on enriched ontologies. Based on the mean expression of the top 1,000 most variable genes (the top 10 genes are shown in Table 2), cell-type centroids were grouped together.

**Table 2.**
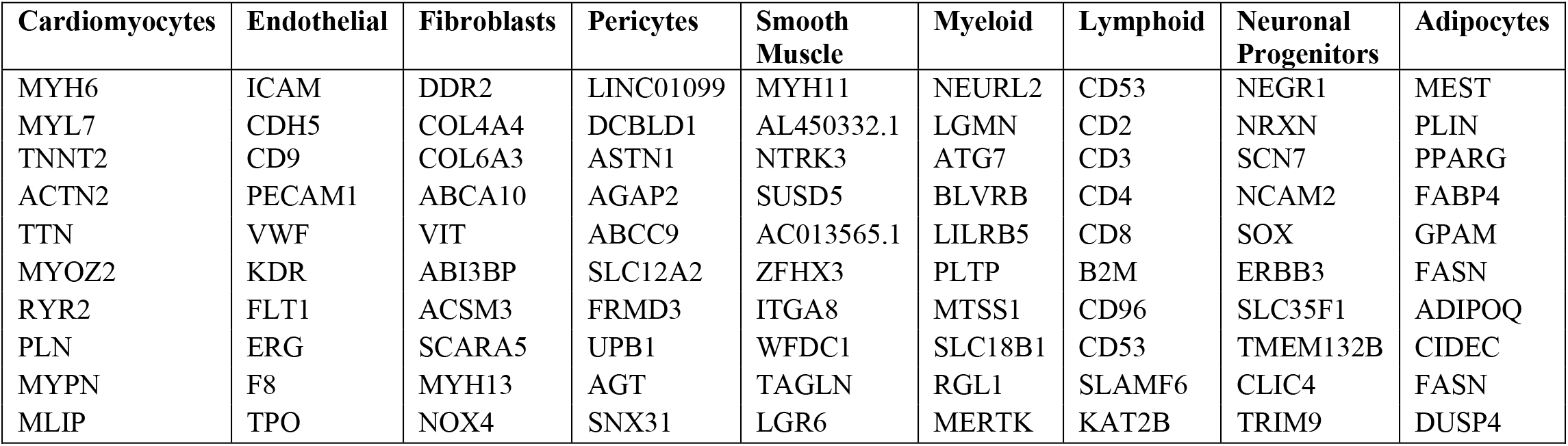
Gene expression profiles of diverse cell types in the heart.

### Single-Nuclei Quality Measure

DNA quality from single cardiac nuclei was measured by multiplex PCR after whole genome amplification^17^. Whole genome amplification was done using Multiple Displacement Amplification (MDA)^18,19^ (Qiagen REPLI-g kit catalog no. 150345) and the BioSkryb Primary Template Amplification (PTA) Kit^20,21^ (Catalog No. PN10013N). Amplification success was determined by Quant-It dsDNA quantification (Thermo Fisher Catalog No. Q33120) in Corning 96-well black absorbance plates (Corning Catalog No. 07-200-590) as well as a multiplexed PCR reaction to assess for even amplification across the genome, with primers targeting regions of chromosomes 5, 10, 15 and 20. Multiplex PCR was performed utilizing 20 ng of DNA with 1x Phusion HF buffer (Thermofisher Catalog No. F549L), 10 mM dNTP, 0.5 U Phusion Hot Start II High Fidelity Polymerase (Thermofisher Catalog No. F549L), and 5-10 μM of the Multiplex Primer Mix (Figure 5e). To further confirm even amplification of each single nucleus, amplified DNA was sequenced (0.5x), and the reads were divided into bins with variable lengths, with each bin having the same number of uniquely mapped reads. Then the differences between copy number ratios of neighboring bins were calculated, and single-nuclei were assigned median absolute pairwise difference (MAPD) scores algorithm^22,23^, where lower values represent even amplification^19^. Higher MAPD scores reflect greater noise. MAPD provides significant advantages over other standard sample deviation measures such as SD, median absolute deviation, and interquartile range. In the present study, based on the observed MAPD score from the 45 single-nuclei, we selected nuclei with MAPD scores of 1.2 or lower for further analysis.

### Cardiomyocyte isolation from frozen tissue and immunostaining for cardiomyocytes and myocardium

Single cardiomyocytes were isolated from the left ventricle of the frozen human heart tissue as described before^24^. Briefly, 100 mg of heart tissue was dissected with a scalpel into small cubes and resuspended in a 1 ml tube containing 500ul of oxygenated cardiomyocyte (CM) isolation buffer containing 130 mM NaCl, 5 mM KCl, 1.2 mM, KH_2_PO_4_, 6 mM HEPES, 1 mM MgCl_2_, 5 mM Glucose, and the sample was swirled gently at room temperature. The buffer was changed to fresh buffer after 3 minutes and the process was repeated twice. Tissue fragments were transferred into a 1 ml tube containing enzymatic buffer (250 μl collagenase (5 mg/ml) and 750 μl CM isolation buffer and 5 μl CaCl_2_) and swirled gently. The sample was incubated while rocking in the enzyme solution at 37°C for 10 minutes, and the digested cells were collected. This process was repeated with undigested tissue until all the cells were dissociated from the tissue blocks. Cells were filtered through a 100 μm filter and centrifuged at 300xg for 2 minutes to pellet the cardiomyocytes. Pelleted cardiomyocytes were resuspended in 1–3 ml (depending on the yield) of 1X PBS. Cells were fixed for 5 minutes, with 100% methanol precooled to −20°C, and the volume was adjusted so that the final concentration was 95%. Fixed cells were centrifuged at 300xg for 5 minutes and the supernatant was discarded. Cardiomyocytes were permeabilized with 0.5% Triton X-100 in 1X PBS at room temperature for 10 minutes and centrifuged at 300xg for 5 minutes, blocked with 1% BSA for 30 minutes at room temperature. The cells were incubated with α-actinin as the primary antibody for 2 hours at room temperature and washed in 1% BSA in PBS twice. Secondary antibody incubation was done for 2 hours at room temperature, and cells were washed twice after incubation with 1% BSA in PBS. For evaluating the myocardium tissue quality, we also performed immunostaining on intact myocardium. The tissue was fixed and embedded in a paraffin block. After deparaffinization, tissues were heated in citrate buffer pH 6.0 (Millipore Sigma) for 20 minutes, permeabilized using 1% donkey serum in PBS plus 0.5% Triton X100, blocked in PBS-T containing 5% donkey serum for 30 minutes at room temperature and incubated with a monoclonal antibody against α-actinin at 1:500 dilution overnight at 4°C. After rinsing in PBST (PBS plus 0.1% Tween-20), sections were incubated with secondary donkey IgG Alexa Fluor 647–conjugated antibody for 1 hour and stained with Wheat Germ Agglutinin and Syto 13 at 1:1000 dilutions for 30 minutes at room temperature. Images were captured on an LSM 880 confocal microscope (Zeiss) and processed using ImageJ (NIH).

### Statistical analysis

Linear regression analysis was conducted to assess the association between DIN or RIN and PMI or age. The Mann-Whitney U test was performed to assess the association between DIN or RIN and gender or race. The Kruskal-Wallis test was performed to assess the association between DIN or RIN and disease status. Scatterplots and boxplots were also created.

## Results

### Quality assessment of heart muscle tissue

We evaluated the tissue quality from 106 collected human heart muscle tissues. The tissues collected were from donors aged .43 to 82 years old, with PMI intervals ranging from 1 to 35 hours, RIN from 4.8 to 9.1, and DIN from 5.8 to 9.8 (Table 1), with higher RIN and DIN values indicating greater nucleic acid integrity. Many of the collected postmortem human heart tissues did not have an associated RIN number when we received the tissue from the NeuroBioBank. For that reason, along with RIN, we evaluated the tissue DNA quality when analyzing sample fit for further single-cell analysis. In our study to evaluate the correlation between RIN, PMI, and age, we performed regression analysis between DIN or RIN and PMI or age in 60 collected heart tissue samples. A Mann-Whitney U test was performed to assess the association between DIN or RIN and gender or race. A Kruskal-Wallis test was performed to assess the association between DIN or RIN and disease status. We found that RIN had no association with age (Figure 2a, p-value=0.140), PMI (Figure 2b, p-value=0.894), race (Figure 2c, p-value=0.937), or gender (Figure 2d, p-value=0.588). Further, we checked for DIN by DNA gel electrophoresis using Genomic DNA Screen Tape. Our study of 37 postmortem fresh frozen heart tissue samples (Table 1a) had an average DIN of 8.39, ranging from 5.8 to 9.8. The DIN value was calculated by the Tapestation 2200 and 4200 by determining the amount of sample degradation present, and DIN values could range from 1 to 10 ^25^. Previous studies have established a DIN of greater than 7 to be an optimal quality of tissue for further biological work^25^. We also tested the DIN from multiple other organs (heart, liver, kidney) for a small number of cases (n=4). Our analysis indicates liver tissue from the same case had a trend of lower DIN with some less than the optimal DIN value of 7 (Table 1b). We found that decreased DIN value was associated with increased age (Figure 2e, p-value=0.007) and increased PMI (Figure 2f, p-value=0.002) using linear regression analysis.

**Figure 2.**
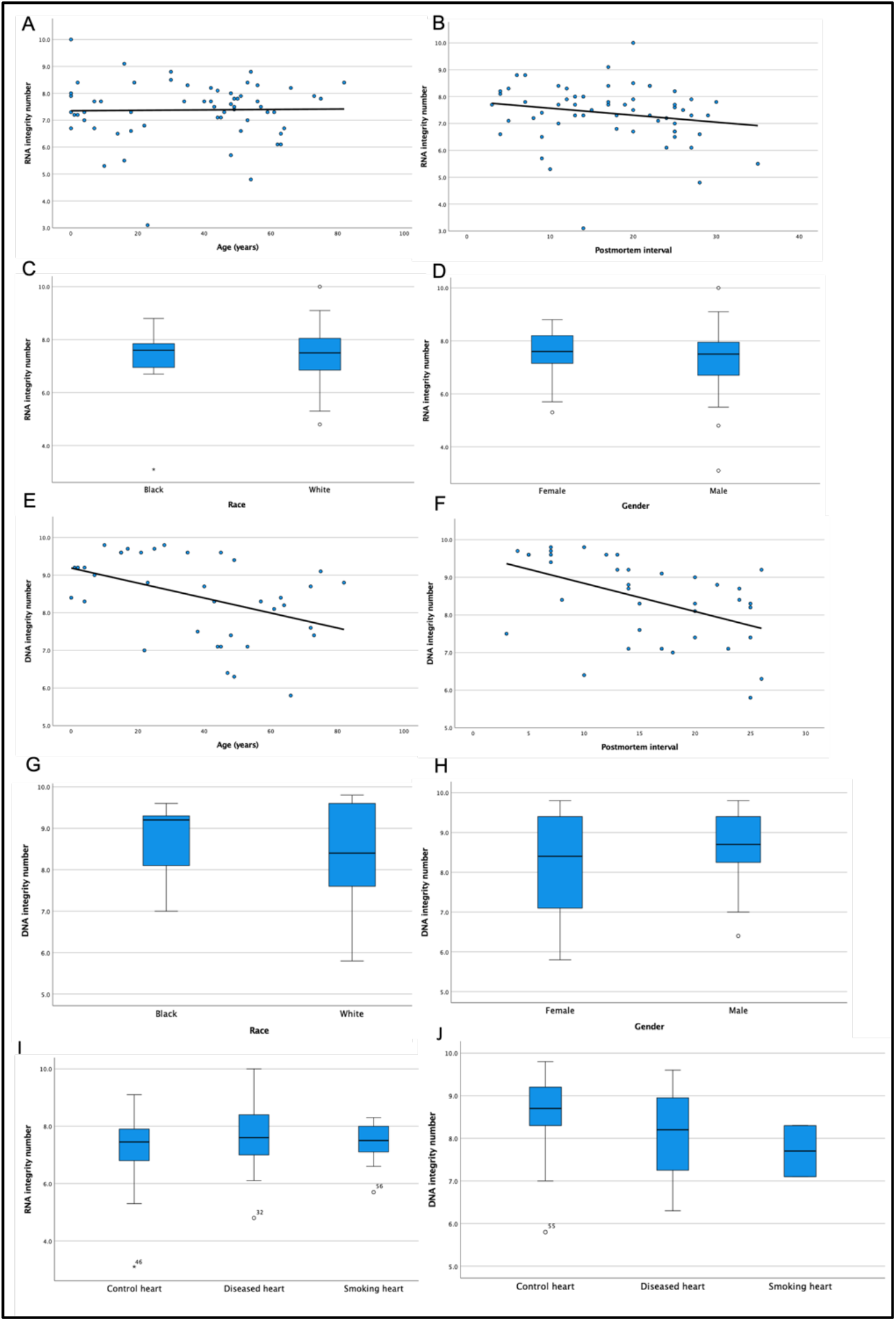
Donor characteristics can impact the quality of heart tissues. (a) Donor age, (b) postmortem interval, (c) race, and (d) gender do not significantly impact the RIN values for heart tissues. However, there is a significant decrease in DIN values of hearts from (e) older donors and (f) from tissues with increased PMI values. (g) Race and (h) gender do not impact DIN values. Disease status does not impact (i) RIN or (j) DIN values.

Analysis of the same sample set for other demographic values such as race and sex showed no association with DIN values for Black/African American versus White/Caucasian donors (Figure 2g, p-value=0.570), or for female versus male donors (Figure 2h, p-value=0.551). Furthermore, we tested the association of RIN and DIN with disease status, characterized as donors with history of atherosclerosis or hypertension as compared to control members; of note, all the DIN values were greater than 7 (Table 1a). We performed the Kruskal–Wallis test for testing the dependence of disease status versus RIN/DIN (Figure 2i, 2j) and found no association (p-value=0.924 and p-value=0.467, respectively). This is notable because many single-cell analysis experiments aim to compare the presence of genome level mutations in control versus disease tissues. If researchers run into problems with their sample preparation, including unsuccessful attempts at whole genome amplification, lack of sufficient amplified sample concentration, or uneven amplification across loci, it is possible that the tissue quality has exacerbated these problems. Therefore, it may be prudent to check the DNA integrity prior to sample preparation to help ensure better results for downstream analysis.

### Determination of quality of cardiomyocytes following single-cell isolation

In this study, the heart tissues we used were collected from the NeuroBioBank. Thus, it was important to evaluate the quality of the heart muscle cells from the tissue bank, where the heart is not the main organ collected in the tissue bank. We tested the quality of isolated cardiomyocytes (Figure 3a, b) from frozen tissue by staining with α-actinin (Millipore-Sigma Catalog No. A7811) as well as a GAP junction protein, connexin 43 (Abcam Catalog No. ab87645). Our immunofluorescence staining with intact connexin in cardiomyocytes (Figure 3c) and α-actinin (Figure 3d) indicated superior quality heart cells could be isolated from these postmortem heart tissues that were collected at the NeuroBioBank. These isolated cardiomyocytes are not for live culture, but they could be used for cell-specific protein expression analysis.

**Figure 3.**
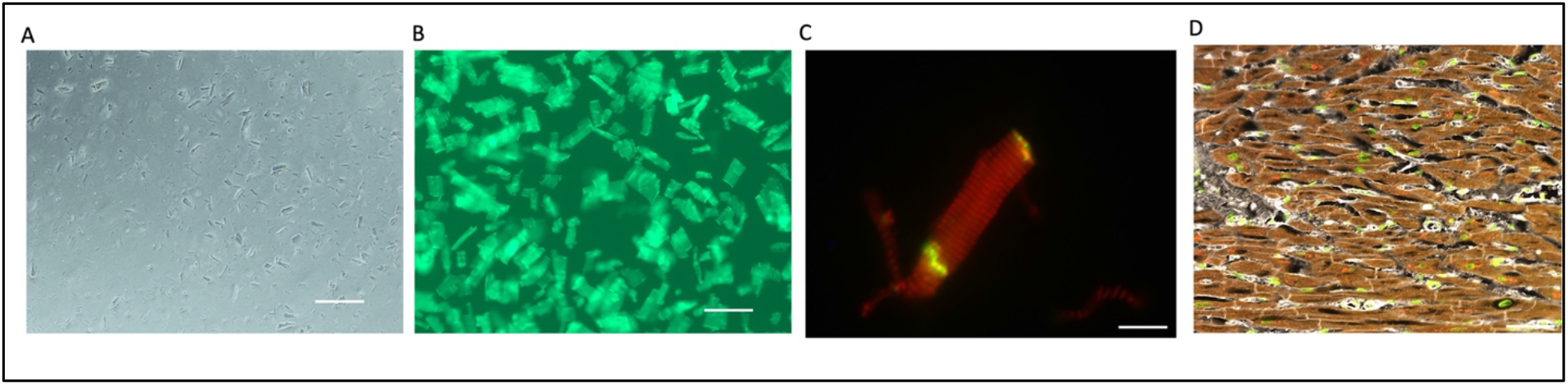
Heart cell and tissue quality assessment. Postmortem heart tissue samples yield high quality single-cell cardiomyocytes. (a) Single-cell cardiomyocytes isolation from postmortem tissue and (b) stained with Cell Brite dye. (c) The single cardiomyocyte was stained with α-actinin (red) and Gap Junction Protein, connexin 43 (green) to determine cell integrity. (d) Heart tissue sections were stained with α-actinin (red) to check for intact myocardium before isolation methods.

### Determination of quality of cardiac nuclei following isolation

To evaluate the ability to isolate and identify all the different heart cell types from these brain-bank-collected heart tissue, we performed 10X Genomics RNAseq (Figure 4a) from a postmortem heart tissue sample with a DIN of 5.8, which was the lowest DIN in our collection, and a heart tissue sample with a DIN of 9.6, which was one of the highest DIN values in our collection (Table 1). We observed an average of 2,000 genes per nucleus and a comparable gene expression profile between these two cases (Supplementary Tables 1 and 2). We identified major cell types in the heart in both samples: cardiomyocytes, fibroblasts (FB), endothelial cells (EC), pericytes, smooth muscle cells (SM), immune cells (myeloid), adipocytes, and neuronal cells (Figure 4b, c). The cell types were identified by the gene expression profiles of the nucleus (Table 2). We also showed the heart cell quality by RNAseq from a heart tissue when the DIN was 9.6 (Figure 4b, Supplementary Table 1) versus when the DIN is 5.8 (Figure 4c, Supplementary Table 2).

**Figure 4.**
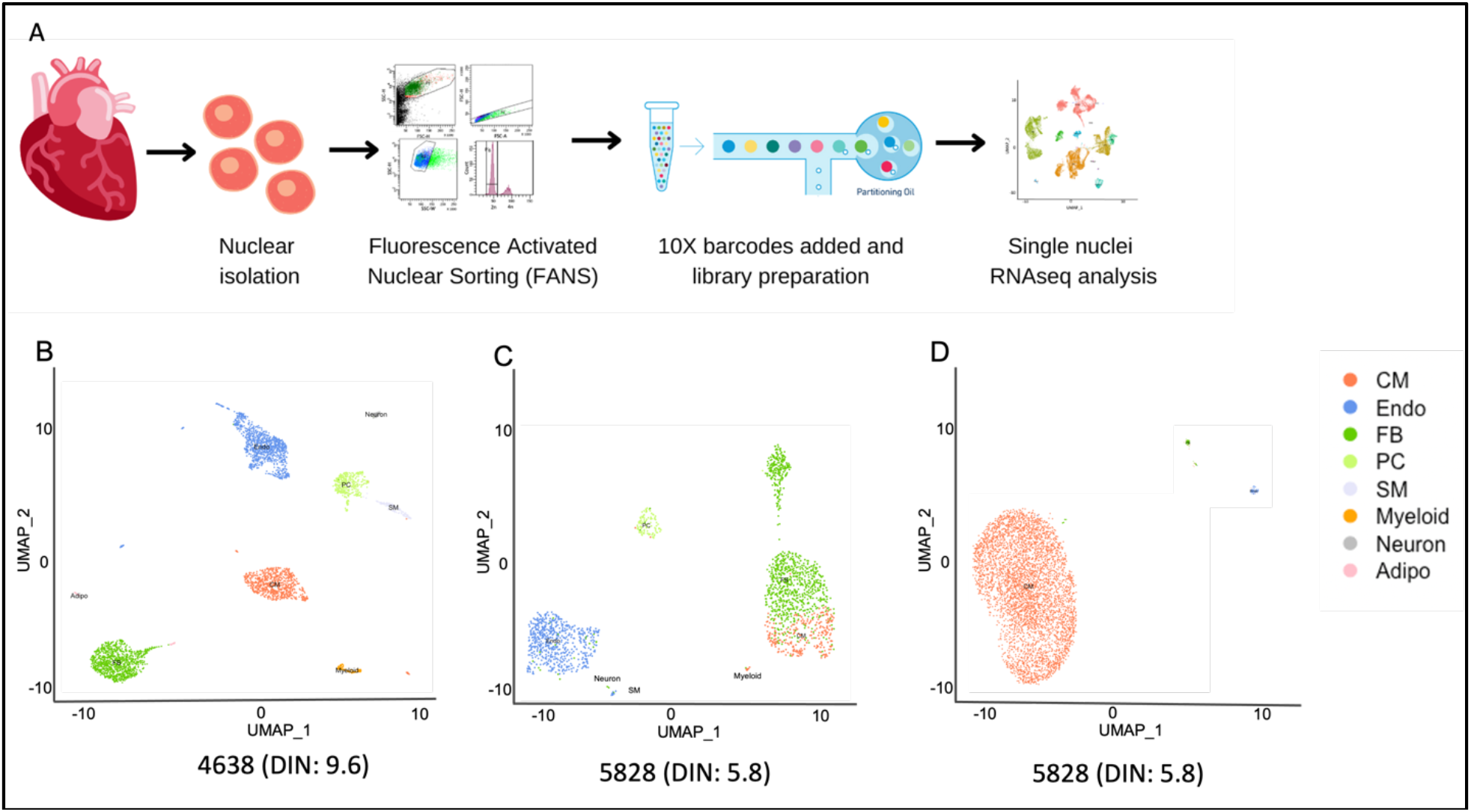
RNAseq data identified all the major cell types of the heart. (a) Schematic of sample preparation for RNAseq. (b) Results of RNAseq data in a case with DIN of 9.6 and (c) a case with DIN of 5.8 in which (d) 95% of tetraploid nuclei in this heart tissue sample were cardiomyocytes with a 3% endothelial cells and 2% fibroblasts. Abbreviations: Cardiomyocytes (CM), Fibroblasts (FB), Endothelial cells (Endo), Pericytes (PC), Smooth Muscle cells (SM), Adipocytes (adipo).

### Determination of DNA quality from single-nuclei isolation

To determine the DNA quality of single-nuclei isolated by the SoNIC method, we performed multiplexed PCR after whole genome amplification by MDA^17,19^ and PTA ^6,21^.

Our analysis indicated successful amplification of all four loci of the selected chromosome locations (Figure 5a, b, c, d) using the primer mix (Figure 5e) in 10-15% of MDA amplified nuclei (Figure 5a, b) and 70-75% in PTA amplified nuclei (Figure 5c, d). The success rate of multiplex PCR was independent of DIN (Figure 5b, d) for our tissue samples which included human tissues with DIN values between 5.8 and 9.8. This finding also indicates that multiplex PCR’s success rate depends on the amplification method. Previous findings have shown improved genome coverage and improved amplification uniformity with PTA^20,21^. Additionally, DIN analysis from multiple organs from the same donor indicated that different organs from the same donor could have a different DIN value (Figure 5f, Table 1b). We also analyzed the median absolute pairwise difference (MAPD) algorithm^22,23^ after a low coverage (0.5x) genome amplification. Although MAPD was originally designed for microarray data, MAPD measures the absolute difference between the log2 copy number ratios of neighboring bins and then calculates the median across all bins. Larger MAPD values indicate greater noise. We found that when the DIN is higher (> 7) we had more amplified cell with MAPD ≤ 1.2, which is considered as the cutoff value for our study (Supplementary Table 3). MAPD analysis indicates that 84.8% of nuclei amplified by MDA had even genome coverage for case 5657 where the DIN was 8.8 and 68.75% for case 5657 where the DIN was 5.8. Together our analysis indicates that SoNIC method could be utilized to isolate superior-quality nuclei from postmortem heart tissue even when the DIN score is 5.8, which is below-average tissue quality.

**Figure 5.**
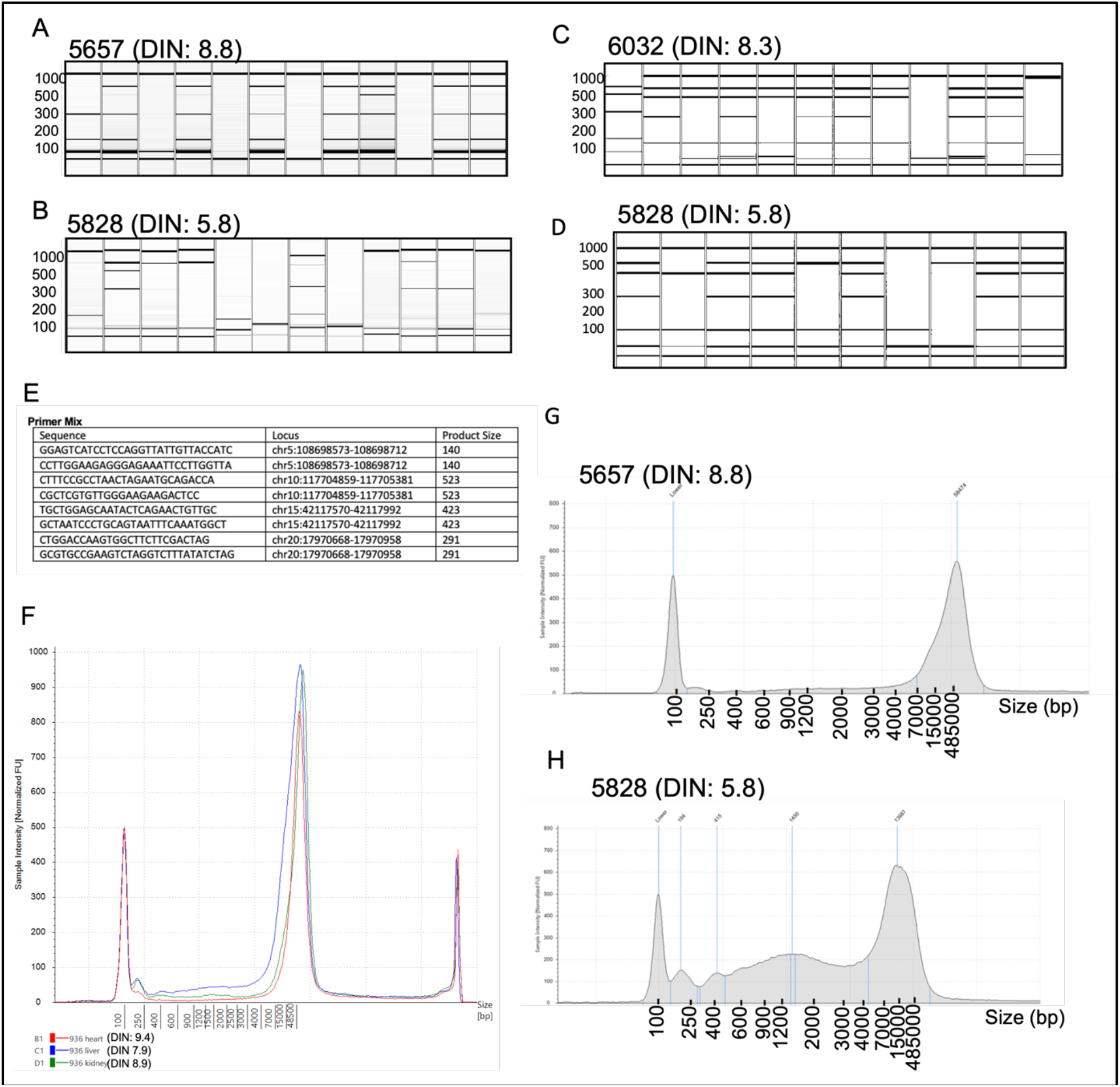
Quality assessment of single cardiomyocyte nuclei and tissue DNA. (a, b) Multiplex PCR product from MDA and (c, d) PTA amplified nuclei are shown on a QIAXcel DNA gel, indicating amplification of chromosome loci. Successful amplification is indicated by the presence of three to four bands. (e) Multiplex PCR primer details with expected product size. (f, g) Tapestation DNA quality analysis on samples with (g) DIN of 8.8 and (h) DIN of 5.8. (f) Comparison of DNA integrity in different tissues from the same human donor.

## Conclusions

Our modified single-nuclei isolation from cardiac tissue (SoNIC) method allowed for the isolation of single cardiomyocyte nuclei from postmortem tissue for single-nuclei whole genome amplification as well as RNAseq. Single-nuclei sorting criteria based on the ploidy of nuclei provided a pure cardiomyocyte nuclei isolation strategy. Furthermore, this study provided detailed quality control steps, summarized in Figure 6 for single-nuclei quality selection criteria for whole genome amplification as well as single-nuclei RNAseq analysis for downstream analysis. In our collected cardiac tissue, we found no association between RIN and PMI, age, or race, whereas the same subset of tissue samples indicated a negative correlation between DIN and PMI or age. Our study emphasizes the inclusion of DIN along with RIN for tissue quality measures prior to the performance of whole genome amplification in human postmortem heart tissue. Our study indicates that postmortem frozen tissue with a DIN over 5.8 could be used for single-nuclei whole genome and RNAseq analysis without compromising the signals important for understanding biological analysis.

**Figure 6.**
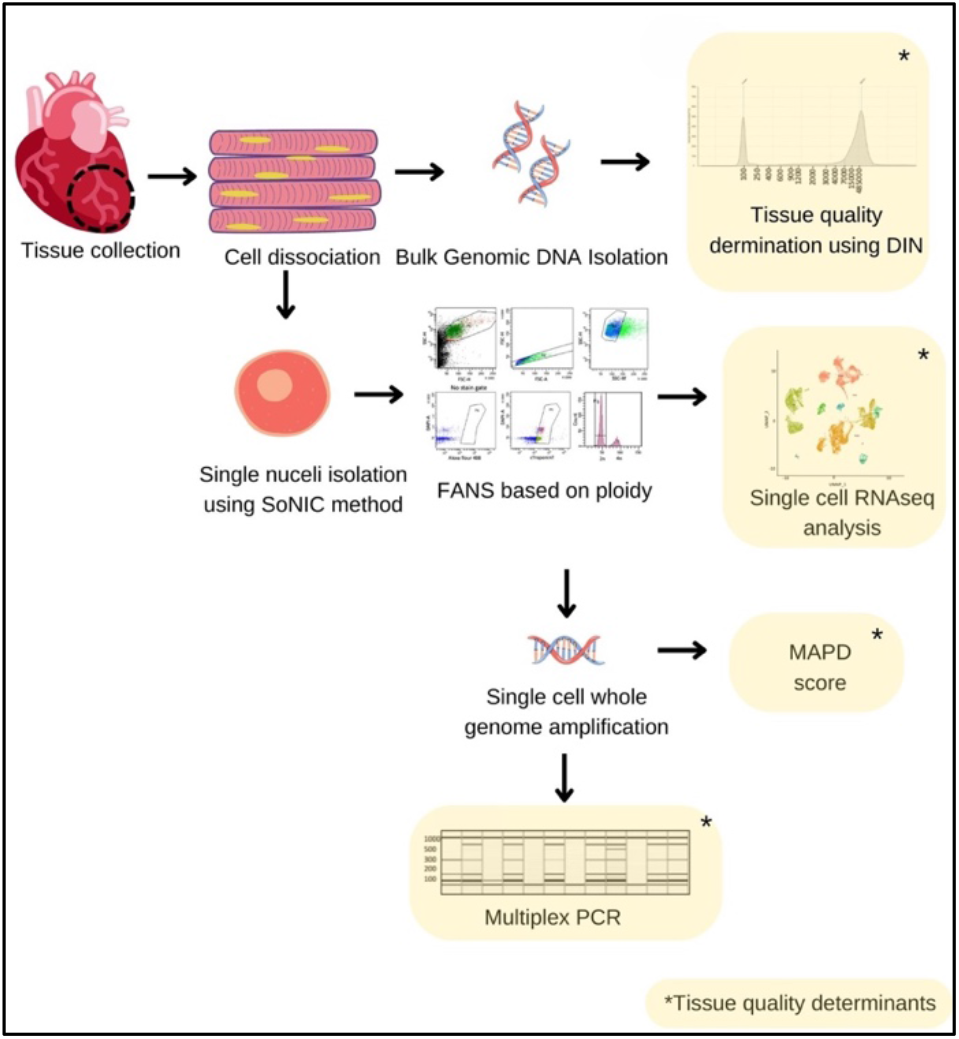
Schematic of approach for tissue quality assessment. Tissue was dissected from the left ventricle of the human heart. Bulk DNA was isolated to check the DNA integrity by Tapestation or the tissue was dissociated to isolate nuclei. After FANS, the single-nuclei were tested for gene expression profile by RNAseq and nuclei were amplified by Φ29 polymerase-mediated MDA for WGS. Successful amplification of single-nuclei was tested via multiplex PCR and genome coverage was evaluated by MAPD score after a low coverage sequencing.

## Limitation of the study

We restricted our analysis to available heart tissue from NIH NeuroBioBank, the University of Maryland (between 2016-2020), which had a DIN value between 5.8 and 9.8.

## Data Availability statement

The original contributions presented in the study are included in the article/Supplementary Material, and further inquiries can be directed to the corresponding author.

## Ethics statement

This study was reviewed and approved by the Boston Children’s Hospital Human Participants Ethics Committee. Postmortem human brain tissue was obtained from the University of Maryland NeuroBioBank. Human tissue was donated there with consent from donors’ families, and its use in this project was approved by the Boston Children’s Hospital IRB.

## Author Contributions

SC contributed to the conceptualization, methodology, analysis, writing, and editing. SA contributed to the methodology, analysis, writing, and editing. RM, AJ, HS, BZ, NH, KM, DN, and IS contributed to the analysis. MHC contributed to the study design, writing, and editing of the manuscript. All authors contributed to the article and approved the submitted version.

## Acknowledgments

We thank R. Sean Hill, Jennifer N. Partlow, Dilenny Gonzalez, and Sonia Epstein for their assistance. Human tissue was obtained from the NIH NeuroBioBank at the University of Maryland, and we thank the donors and their families for their invaluable donations for the advancement of science. SC is supported by American Heart Association (SDG) and NHLBI (R01HL152063). MHC is supported by NHLBI (R01HL152063). AJ was supported by PRISE and HCRP Fellowship Awards from Harvard College. HS was supported by HCRP Fellowship Award from Harvard College.

